# Phylogeny-aware modeling uncovers molecular functional convergences associated with complex multicellularity in Eukarya

**DOI:** 10.1101/2024.03.14.585020

**Authors:** Francisco Pereira Lobo, Dalbert Benjamim da Costa, Thieres Tayroni Martins da Silva, Maycon Douglas de Oliveira

**Affiliations:** Laboratório de Algoritmos em Biologia, Departamento de Genética, Ecologia e Evolução, Instituto de Ciências Biológicas, Universidade Federal de Minas Gerais, Brazil

**Keywords:** Evolution of Multicellularity, Molecular Functional Convergence, Homology, Genotype-Phenotype Association, Comparative Genomics

## Abstract

A major trait of Eukarya is the independent evolution of complex multicellular lineages of animals and plants with specialized cell types organized in tissues, organs and systems. The number of cell types (NCT) has been commonly adopted as a proxy in comparative studies investigating the genomic evolution of biological complexity. Although expansions of homologous genes playing roles in extracellular processes, signal transduction pathways, and the immune system have been reported as associated with NCT variation in metazoans, the evolutionary patterns coordinating the genomic evolution of multicellularity throughout Eukarya remain poorly understood. We used Gene Ontology (GO) as a genome annotation scaffold to represent biological functions shared by genes regardless of their homology relationships to search for molecular functional convergences associated with NCT. For that we integrated phenotypic, genomic, annotation and phylogenetic data to build phylogeny-aware models and searched for expansions of homologous regions and of GO terms associated with NCT values across 49 eukaryotic species, including complex multicellular plants and metazoans. Virtually all homologous regions associated with NCT are metazoan- and vertebrate-specific expansions of paralogs with key roles in developmental pathways. The functional annotation, in contrast, detected previously unknown biological themes coded by non-homologous genes independently expanded in multicellular plants and metazoans, such as system and anatomical development, immunity, regulatory mechanisms of embryogenesis, response to external stimuli and detection of natural rhythmic processes. Our findings unveil functional molecular convergences shared by these contrasting groups due to common selective pressures in the lifestyle of complex multicellular lineages.

## Introduction

A core trait in the evolution of Eukarya is the multiple emergence of multicellular lineages, such as fungi, animals and plants. These are defined as taxa whose somatic bodies contain specialized cell types with cell-cell communication mechanisms and task division. Additionally, complex multicellularity evolved independently in certain lineages of metazoans and embryophytes (land plants) with the emergence of a hierarchical organization of cells in tissues, organs and systems^1,2^. The independent evolution of multicellularity in Eukarya suggests that this taxon possesses a set of shared homologous sequences that contributes for the multiple appearance of this phenotype^3^. It is also reasonable to assume that some of the molecular components associated with the independent emergence of multicellularity are molecular functional convergences, where non-homologous genes fulfill common biological roles in distinct multicellular eukaryotic lineages. A compelling example is the independent expansion of non-homologous developmental transcription factors in multicellular plants and metazoans^1^. Therefore, the search for molecular functional convergences associated with complex multicellularity can benefit from annotation schemas that reflect the functional similarities of non-homologous genes.

Gene Ontology (GO) is a controlled dictionary of annotation terms and their relationships that describes the biological properties of gene products regardless of their homology relationships^4^. GO terms have been widely adopted to help interpreting large-scale omics experiments through the search of common biological themes in gene lists^5^. However, the adoption of GO as a function-based gene annotation scaffold for comparative genomics studies remains limited, despite its ability to capture convergent molecular patterns in the evolution of complex phenotypes, such as parasitism and sociality^6,7^.

The number of cell types (NCT) has been widely adopted as a proxy to quantify the biological complexity across eukaryotic species in comparative genomics studies to understand the molecular evolution of this phenotype^8–11^. NCT variation is positively associated with many properties that quantify genomic complexity in some degree, such as the size of non-redundant proteomes and the number of alternative splicing events^9,10^. Nevertheless, these general associations do not inform about specific groups of homologs and molecular functional convergences are associated with NCT variation. Although expansions of homologous genes from extracellular processes, signal transduction pathways and components of the immune system are associated with NCT variation in metazoans^11^, the evolutionary patterns coordinating the genomic evolution of multicellularity throughout Eukarya remain poorly understood.

Furthermore, the development of statistical workflows for exploring genotype-phenotype associations across species sharing common ancestors requires one to consider data interdependence caused by shared ancestry^12^. Here we report the large-scale integration of phenotypic, genome annotation and phylogenetic data to build phylogeny-aware linear models aimed at searching for homologous regions and functional convergences associated with NCT variation. For that we evaluated 49 phylogenetically diverse eukaryotic species with varying degrees of complexity including multicellular plants and metazoans, the two eukaryotic lineages where complex multicellularity emerged independently.

We demonstrate how annotations as provided by GO terms outperform homology-based ones by offering a significantly higher set annotation terms shared across the genomes of multicellular organisms that are, consequently, more suitable for detecting functional molecular convergences. Homologous regions associated with NCT correspond virtually to metazoan-specific expansions of gene families playing key roles in cell identity, embryogenesis, organogenesis, and system development, together with poorly characterized gene families that are interesting targets for experimental characterization. In contrast, GO annotation unveiled many previously unknown, independent expansions of biological themes in multicellular plants and animals. These include concepts long though as essential for the emergence and maintenance of complex multicellularity, such as developmental processes and regulation of cell proliferation. We also report new and unexpected biological roles independently expanded in the genomes of these lineages, such as the detection of circadian rhythmic processes, the response to wounding, and the response to external stimuli.

This study showcases the effectiveness of utilizing GO terms and phylogenetic comparative methods to identify molecular convergences in comparative genomics investigations. More importantly, we demonstrated how complex multicellular organisms have independently evolved functionally similar gene sets to interact with their dynamic environments throughout their life cycles.

## Results

### Experimental design

We integrated phylogenetic, genome annotation, and phenotypic data to build linear models and search for associations between NCT and genes annotated to reflect either their homology or functional relationships (Supplementary Table 1; Figure 1A; Methods, section “Acquiring high-quality phylogenetic, annotation and phenotypic data”). For that purpose we use CALANGO^13^, a phylogeny-aware comparative genomics tool that searches for quantitative genotype-phenotype associations, flagging as positive the associations with q-values < 0.01.

**FIGURE 1.**
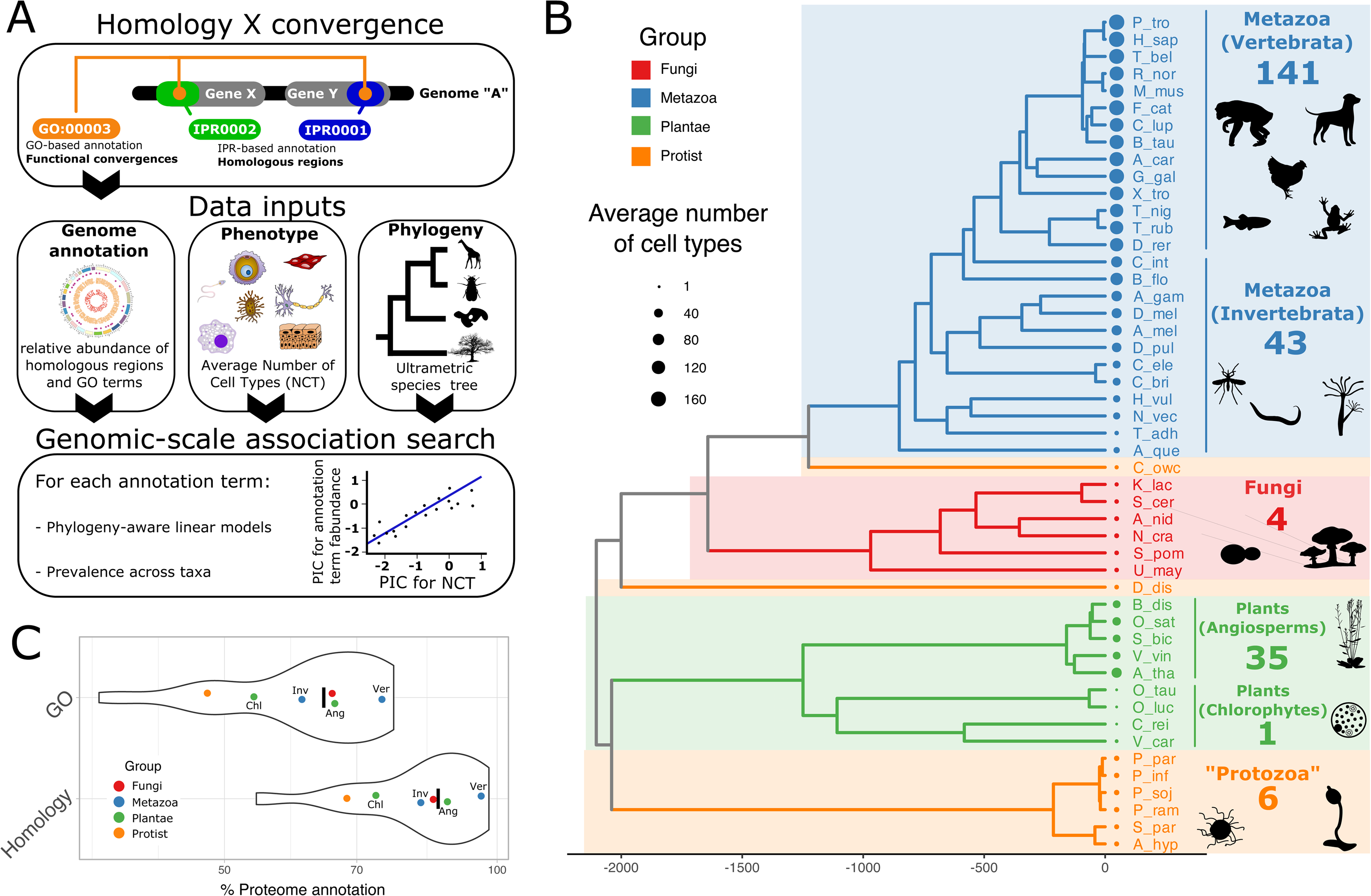
Experimental design, phenotypic, phylogenetic and genomic data. A) Experimental design. In brief, our strategy relies on the fact that non-homologous regions that fulfill the same biological roles (green and blue boxes) will be annotated to the same GO term (“Homology versus convergence”). We integrated genome annotation, phenotypic and phylogenetic data to build phylogeny-aware models and search for homologous regions and biological roles associated with the number of cell types (NCT) across Eukarya. **B)** NCT values across major eukaryotic lineages. Species acronyms are as follows: A_car (*Anolis carolinensis*), A_gam (*Anopheles gambiae*), A_hyp (*Achlya hypogyna*), A_mel (*Apis mellifera*), A_nid (*Aspergillus nidulans*), A_que (*Amphimedon queenslandica*), A_tha (*Arabidopsis thaliana*), B_dis (*Brachypodium distachyon*), B_flo (*Branchiostoma floridae*), B_tau (*Bos taurus*), C_bri (*Caenorhabditis briggsae*), C_ele (*Caenorhabditis elegans*), C_int (*Ciona intestinalis*), C_lup (*Canis lupus*), C_owc (*Capsaspora owczarzaki*), C_rei (*Chlamydomonas reinhardtii*), D_dis (*Dictyostelium discoideum*), D_mel (*Drosophila melanogaster*), D_pul (*Daphnia pulex*), D_rer (*Danio rerio*), E_cun (*Encephalitozoon cuniculi*), E_his (*Entamoeba histolytica*), F_cat (*Felis catus*), G_gal (*Gallus gallus*), H_sap (*Homo sapiens*), H_vul (*Hydra vulgaris*), K_lac (*Kluyveromyces lactis*), L_maj (*Leishmania major*), M_mus (*Mus musculus*), N_cra (*Neurospora crassa*), N_vec (*Nematostella vectensis*), O_luc (*Ostreococcus lucimarinus*), O_sat (*Oryza sativa*), O_tau (*Ostreococcus tauri*), P_bal (*Populus balsamifera*), P_fal (*Plasmodium falciparum*), P_inf (*Phytophthora infestans*), P_par (*Phytophthora parasitica*), P_pat (*Physcomitrella patens*), P_soj (*Phytophthora sojae*), P_tro (*Pan troglodytes*), R_nor (*Rattus norvegicus*), S_bic (*Sorghum bicolor*), S_cer (*Saccharomyces cerevisiae*), S_moe (*Selaginella moellendorffii*), S_par (*Saprolegnia parasitica*), S_pom (*Schizosaccharomyces pombe*), T_adh (*Trichoplax adhaerens*), T_ann (*Theileria annulata*), T_bru (*Trypanosoma brucei*), T_gon (*Toxoplasma gondii*), T_nig (*Tetraodon nigroviridis*), T_rub (*Takifugu rubripes*), U_may (*Ustilago maydis*), V_car (*Volvox carteri*), V_vin (*Vitis vinifera*), X_tro (*Xenopus tropicalis*), Y_lip (*Yarrowia lipolytica*), Z_may (*Zea mays*). **C)** Genome annotation coverage (percentage of protein-coding genes associated with at least one annotation term) for GO-based and Homology-based schemas. Acronyms are defined as follows: Vert – Vertebrates; Invert – Invertebrates; Angio – Angiosperms; Chlor – Chlorophytes.

The phenotypic data comprises average NCT values for each species gathered from the literature with the exception of *Homo sapiens*, for whom we used the data from *Pan troglodytes* (common chimpanzee) as a proxy. This was necessary since NCT values for our species are inflated due to its larger amounts of single-cell data (Supplementary File 1, section “NCT value of *Homo sapiens* is biased when compared with other species and influences downstream analyses”). The genomic data consists of 49 high-quality non-redundant proteomes annotated using either homology-based (InterPro IDs, from now on referred as IPRs) or function-based annotation schemas (GO IDs) (Supplementary Figures 1A-B). By annotating the same set of components to these two annotation schemas, we could also objectively compare the distinct annotation strategies for their capabilities to detect molecular convergences. CALANGO also requires a species tree to compute phylogenetically independent contrasts for both the annotation and phenotypic data. We integrated phylogenetic data from two trees: one scaffold tree representing the most accepted early branching patterns of eukaryotic evolution^14^ and one donor tree containing the phylogenetic information for more recent groups^15^. We also incorporated recent information from early metazoan evolution to reflect its evolutionary history^16^ (Methods, section “Phylogenetic data”; Supplementary Figures 1C-E).

### Evolution of NCT in *Eukarya*

Metazoans and multicellular plants have the highest NCT values, while Fungi have a mean NCT value similar to basal unicellular eukaryotic species with complex life cycles and distinct cell types as previously reported^1,11,17^ (Figure 1B). Within metazoans, vertebrates have a substantial increase in NCT, with an average of 141 cell types compared to an average NCT value of 43 for invertebrates. The same qualitative pattern is observed within Archaeplastida, where chlorophytes have some of the smallest NCT values (average of one), whereas angiosperms have much higher number of cell types (average of 35). In addition, invertebrates have NCT values similar to multicellular plants. Together, as we demonstrate below, these independent events of increase of NCT in multicellular plants and metazoans allow the search for independent expansions of both homologous regions and functional roles associated with these NCT expansions.

### Properties of homology-based and functional-based annotation schemas

We built two annotation schemas to compare traditional homology-based annotation schemas with functional, GO-based annotations. In brief, we generated unique pairs of proteins and their annotation terms, when available, representing gene-level counts of either homologous regions (IPR IDs) or GO terms for each genome (Supplementary Figure 1A). The homology-based annotation consisted of 2,685,970 unique pairs of 741,033 distinct gene products annotated to at least one out of the 24,502 distinct IPR IDs from our *de novo* annotation. As for the GO-based annotation, we found 1,490,781 unique pairs of 555,811 gene products annotated 8,820 distinct GO terms.

The percentage of proteome annotation on each database, defined as the fraction of proteins annotated to at least one annotation term from that database, is highly variable in our dataset. The GO-based schema annotates a smaller fraction of the proteome than its homology-based counterpart (Figure 1C). This is due to the several conserved regions formally described by entries in the homology-based databases that nevertheless lack deeper functional characterization and, consequently, have not been associated to any GO IDs.

The percentage of proteome annotation is also biased towards taxa containing economically relevant model organisms. Vertebrates have the largest proteome annotation coverage, with angiosperms, invertebrates and fungi having intermediate values, and chlorophytes and protozoans have the lowest coverage values. This observation was previously reported^18^ and reflects our lack of biological knowledge about most of the gene products from neglected taxa. Another source of variation in the counts of annotation terms is the considerable differences in the size of non-redundant proteomes in Eukarya, again with metazoans and plants having the highest values (Supplementary Figure 1B).

Together, these facts are likely to introduce biases in the estimation of the abundance of annotation terms across genomes and in our downstream analyses. We accounted for this by using the total number of unique pairs in a genome as a correction factor to compute the relative frequencies of annotation pairs in each genome. In this scenario, vertebrate proteomes, for instance, have the largest correction factors, as these species have both the largest fraction of annotated proteomes and also the largest non-redundant proteomes, and both factors contribute for increase of annotation pairs.

We also assessed the prevalence of each annotation term, defined as the fraction of genomes where each annotation term was observed. Since annotation terms with higher prevalence are observed in more genomes and, therefore, are more taxonomically diverse, they are also more useful to detect independent expansions of annotation pairs in phylogenetically distant lineages (Figure 1A, Supplementary Figure 1A). To investigate the prevalence of annotation terms across the distinct multicellular eukaryotes, we filtered out protist-exclusive terms (around 1% of all annotation terms; 272 out of 24,502 homologous regions, for instance) and classified the remaining groups as either lineage-specific (when found exclusively in a single multicellular lineage) or shared (when found in two or more multicellular lineages) (Figure 2).

**FIGURE 2.**
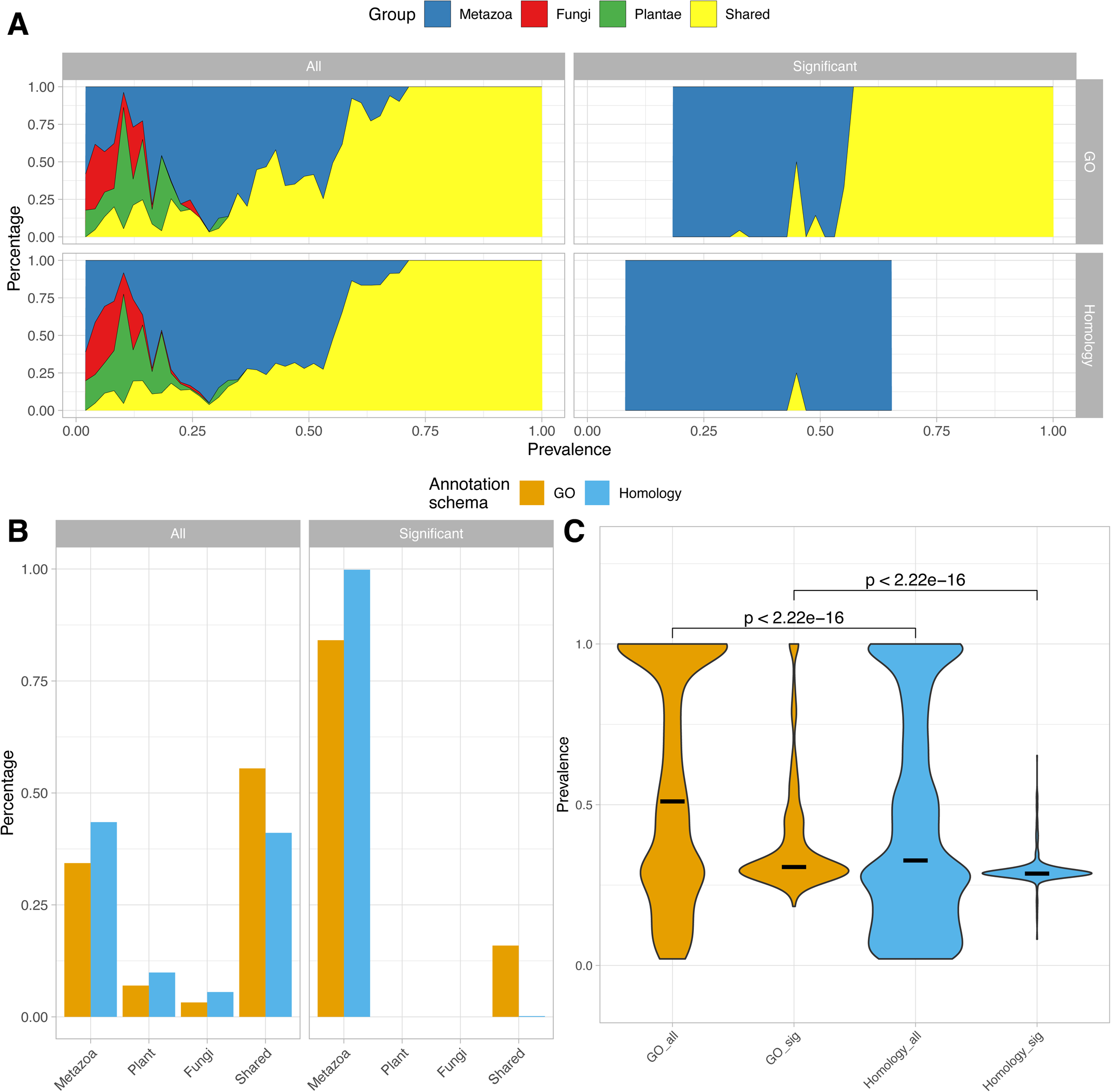
Comparison of function- and homology-based annotations for the detection of molecular convergences. A) Taxonomic abundance of annotation terms with varying prevalence values for the distinct annotation schemas (prevalence defined as the fraction of genomes where a given annotation term was observed). Data in panels is as follows: “All” – full set of annotation terms, “Significant” – annotation terms associated with NCT, “GO” – Gene Ontology annotation terms, “Homology” – InterPro annotation terms. B) Fraction of annotation terms exclusively observed in one multicellular lineage (Fungi/Metazoa/Plantae) or shared across at least two multicellular taxa. C) Prevalence distribution across genomes for different annotation schemas. Groups are defined as follows: “GO_all” – all annotation terms from the GO-based annotation schema, “GO_sig” – GO-based annotation terms associated with NCT, “Homology_all” – all annotation terms from the homology-based annotation schema, “Homology_sig” – homology-based annotation terms associated with NCT.

We found a similar taxonomic distribution of annotation terms as a function of their prevalence in the two annotation schemas. Terms with lower prevalence are mostly lineage-specific, with metazoan-specific annotation terms being the most abundant. This is especially true for prevalence values around 0.3, which corresponds to the fraction of vertebrate genomes in our dataset (Figure 2A, “All” column, metazoan-specific terms represented as blue polygons). As expected, terms with higher prevalence are shared across at least two multicellular eukaryotic lineages (Figure 2A, “All” column, yellow curves).

Shared terms are the largest category in both annotation schemas, with metazoan-specific groups being the second largest group (Figure 2B), yet another evidence of the abundance of functional characterization of genes from this taxon (Figure 1C). However, the homology-based annotation had a higher fraction of lineage-specific terms compared to the GO-based annotation, as the later contains a higher fraction of shared terms. The higher prevalence of GO terms indicates that the function-based annotation is annotating non-homologous genes fulfilling the same biological roles to common GO IDs, making them more suitable for the detection of molecular convergences.

The distribution of prevalence for homologous regions is bimodal, with a smaller peak of highly prevalent terms and a larger peak with smaller prevalence values. The majority of values around 0.3 corresponds to the peak of annotation terms from vertebrate genomes previously observed (Figure 2C, “Homology_all”). The distribution of prevalence for GO terms is also bimodal, but with a significantly larger fraction of terms with higher prevalence values (Figure 2C, Wilcoxon test, “GO_all” vs “Homology_all”). This provides additional evidence that non-homologous regions that fulfill the same biological roles have been annotated to common GO terms.

### GO terms associated with NCT are significantly more prevalent across eukaryotic genomes compared to associated homologous regions

We found 607 homologous regions associated with NCT (2.47% of the total number of 24,502 entries of the homology-based annotation). As for the functional-based annotation, we found 352 associated GO terms (3,99% of the 8,820 GO terms observed in at least one genome) (Supplementary Table 2 contains all annotation terms deemed as associated for the two annotation schemas). The GO-based annotation schema generated associations with higher prevalence values compared with the homology-based annotation and, consequently, shared across multicellular lineages (Figure 2A, “Significant” column).

Even though the largest fraction of annotation terms are shared across one or more multicellular lineage regardless of the annotation schema (Figure 2B, “All”), virtually all of the homologous regions associated with NCT values are exclusively observed in metazoans (Figure 2B, “Significant” column). In fact, only one out of the 607 (0.16%) distinct homologous regions associated with NCT is shared across multicellular lineages. In contrast, 15,9% (56 out of 352 terms) of the GO terms associated with NCT values also occur in more than one multicellular lineage (Figure 2B, “Significant” panel). The prevalence of GO terms associated with NCT is significantly higher than the homology-based associations (Figure 2A, “GO_sig” vs “Homology_sig”). Both distributions have the sharp peak around 0.3 corresponding to vertebrate-specific expansions, but only the list of associated GO terms have a long tail of terms with higher prevalence.

Together, these observations demonstrate how the GO-based annotation schema, when compared to the homology-based annotation, recovers a higher fraction of terms associated with NCT that are also shared by distinct multicellular taxa. We conclude that GO-based annotation schemas are likely to be detecting molecular functional convergences not found by the homology-based annotation schema.

### Homologous regions associated with NCT comprise metazoan- and vertebrate-specific expansions of key genetic components of multicellularity

All but four of the 607 IPR terms associated with NCT correspond to metazoan-specific expansions, with 78,3% of the associations restricted to the vertebrate lineage (475 homologous regions) (Figure 3A, “IPR” heatmap; see also Supplementary File 1, section “Analysis of homologous regions associated with NCT and broader taxonomic distribution”). The single IPR domain associated with NCT and shared by two distinct lineages where multicellularity evolved is the P2X purinoceptor (IPR001429, Supplementary Figure 3A). This entry was observed in metazoans and in the chlorophytes *Ostreococcus tauri* and *O. lucimarinus* (Archaeplastida), some of the organisms with the smallest cell type counts in our dataset (Figure 1B). This entry is also observed in *Capsaspora owczarzaki* (Opisthokonta), with a clear expansion in metazoans, but is absent from the other eukaryotic multicellular lineages (angiosperms and fungi)^19,20^. In metazoans, P2X receptors are involved in neurotransmission, neuromodulation, and immunological roles, with no known roles for this gene in chlorophytes^19,20^. Overall, we conclude that this homologous region is likely to play an important role for the emergence of multicellularity in metazoans. However, it clearly does not comprise compelling evidence of an independent expansion in multicellular eukaryotes.

**FIGURE 3.**
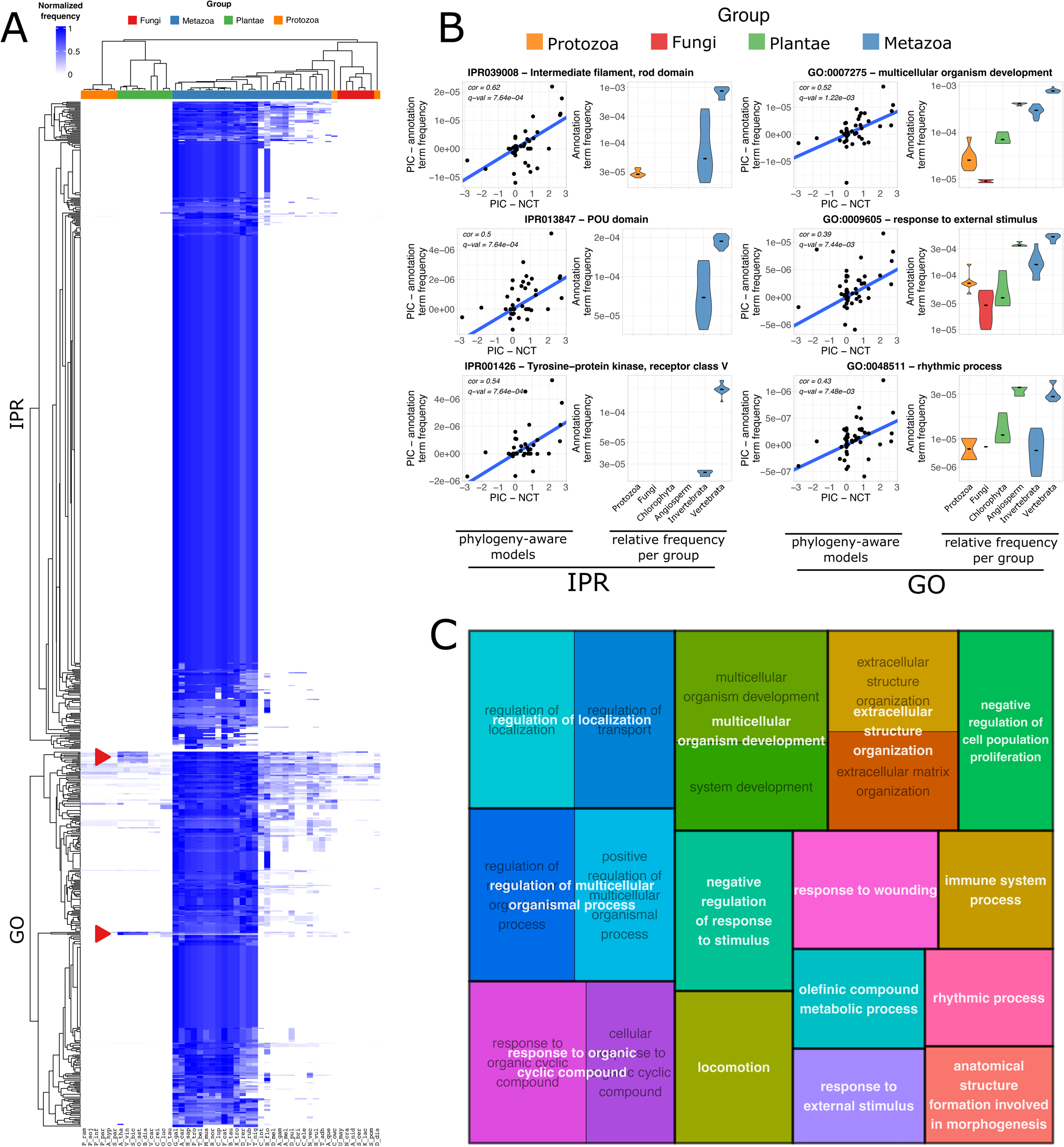
homologous regions and functional roles associated with the evolution of multicellularity in Eukarya. A) Heatmaps of IPR IDs and GO term IDs associated with NCT. Species codes are the same as in Figure 1A. Red arrows highlight independent expansions of GO terms in plants and metazoans. B) Examples of homologous regions and GO terms associated with NCT values. Left: association between phylogenetically independent contrasts (PIC) for NCT and for the relative frequency of annotation terms. The blue line represents the linear model of PICs; Correlation values are Pearson’s correlation of PIC values; q-values are corrected p-values for the linear models. Right: relative frequency of the annotation terms for the six groups investigated in this analysis. Each violin plot represents the distribution of values for the group; black bars are the medians of group. C) Treemap of GO terms associated with NCT values independently expanded in angiosperms and metazoans. Each box is proportional to the corrected q-values of the phylogeny-aware linear models.

The other three homologous regions associated with NCT observed outside metazoans are exclusively present in this lineage and in eukaryotic unicellular species within the protozoan group. These were selected for further inspection, as they represent pre-metazoan homologous regions that may have contributed for the emergence of multicellularity in this lineage. Overall, we found our results consistent with the known taxonomic distribution of these gene families (Supplementary File 1, section “Analysis of homologous regions associated with NCT and broader taxonomic distribution”). These homologs have been previously reported to have a pre-metazoan origin and correspond to expansions of gene families previously described as involved in the evolution of multicellularity, such as intermediate filaments and extracellular matrix metalloproteinases. These genes play roles in cell behaviors such as proliferation, migration, differentiation, and determination of cell shape^21,22^ (Figure 3B, “Homology Column”; Supplementary Figures 3B-C). Even though these domains are clearly expanded in metazoans and associated with the evolution of multicellularity in this lineage, they fail to be classified as independent expansions in distinct eukaryotic multicellular lineages.

All remaining 603 groups of homologs associated with NCT are exclusively observed in metazoans (99,3% of the associated regions). From those, 132 associated homologous regions were observed in invertebrates and vertebrates. These include gene families previously reported as metazoan expansions that are key components of the extracellular matrix and mediators of cell-cell interactions, such as Syndecans (IPR001050) and Laminins (IPR009254)^23–25^ (Supplementary Figures 4A-B). Other expansions correspond to gene families playing distinct roles in developmental processes, such as ion channels (IPR002946 – *Intracellular chloride channel*)^26^, hormone receptors (IPR001426 – *Tyrosine-protein kinase, receptor class V, conserved site*)^24^, transcription factors (IPR013847 – *POU domain,* IPR015395 - *C-myb, C-terminal*)^27^, epigenetic regulators (IPR016376 – *Histone acetyltransferase GCN5/PCAF*)^28^, and caspases (IPR015471 − *Caspase−3/7*)^29^ (Figure 3B, Supplementary Figures 4C-H). Other expansions included components of canonical developmental pathways, such as Wnt and Homeobox (IPR016233 - *Homeobox protein Pitx/unc30*; IPR013301 - *Wnt-8 protein, Supplementary*; Figures 5A-B)^30^, together with tissue-specific growth factors (Supplementary Figures 5C-E). Many metazoan-restricted associations are poorly characterized groups of homologs that constitute interesting targets for experimental characterization in the search for new components that may have contributed for the evolution of multicellularity in this lineage (Supplementary Figures 5 F-J).

The remaining 475 associations are vertebrate-specific expansions that correspond mostly to the same biological roles previously described in metazoans, including proteins of the extracellular matrix, hormones, components of developmental signaling pathways and poorly characterized gene families (Supplementary Figures 6 A-L). Together, our findings present compelling evidence that the homology-based annotation schema detected expansions of many genes families that are fundamental drivers of multicellularity in metazoans. This annotation schema, however, failed to find any homologous region independently expanded in eukaryotic multicellular lineages.

### Comparison of our work to previous studies

To the best of our knowledge, a single study by Vogel and Chothia (from now on referred as VC work) is the only work to search for expansions of specific protein families associated with NCT variation in Eukarya^11^. These authors found a majority of metazoan-specific gene expansions enriched in genes playing roles in extracellular processes, signal transduction and components of the immune system. However, using traditional correlation statistics to detect associations in cellular organisms is unsound due to non-independence of data of species caused by common ancestry^12^. Therefore, we investigated how applying our phylogeny-aware models affects their findings.

A fraction of the homology-based annotation terms used in this study are defined by the SUPERFAMILY database, one of 13 databases providing annotation terms to InterProScan. This is also the same annotation schema used by the VC work. Thus, the SUPERFAMILY annotation schema provides an interesting point of comparison to analyze the findings and conclusions of both works. We initially employed a statistical model to emulate their statistical strategy by do not accounting for the non-independence of species data when searching for associations. Our results using these models significantly agreed with theirs, indicating that the differences between our datasets, such as the greater number of species, the updated NCT values, and the newer genome versions in our dataset do not influence the overall conclusions of VC (Supplementary File 1, section “Failing to take into account phylogenetic relatedness produces spurious associations”). However, when we employed the phylogeny-aware models to search for such associations, no homologous regions nor GO terms in the SUPERFAMILY annotation schema were found to be significantly associated with NCT values.

### Molecular functional convergences independently expanded in multicellular lineages

The 56 associated GO terms associated with NCT and observed in at least two of the three multicellular lineages were evaluated for molecular convergences by searching for independent expansions in metazoans and other multicellular lineages. For that, we searched for annotation terms where either fungi or angiosperms where among the three highest non-zero median values across the six groups represented in Figure 1B (see “Methods”, section “Detection of molecular functional convergences”). This analysis labelled 26 out of the 56 shared GO terms as metazoan-specific expansions. These correspond to terms with a higher relative abundance in the two metazoan groups compared with the values in the other multicellular lineages, whose values were comparable to groups of unicellular/oligocellular eukaryotic species (e.g. GO:0016247 – *channel regulator activity*, GO:0009968 – *negative regulation of signal transduction*; Supplementary Figures 7A-B). Even though these GOs correspond to ancient biological processes whose individual components have existed since before the emergence of multicellularity and contributed for the emergence of multicellularity in metazoans, they do not provide evidence of molecular functional convergences.

Our strategy to find molecular convergences labelled 30 GO terms as independent expansions in the relative abundance of key biological functions for the evolution of multicellularity in at least two multicellular lineages. Eight of these GO terms are independently expanded in Fungi and metazoans. Among these terms we highlight GO:0007265 – *Ras protein signal transduction*, independently expanded in metazoans and fungi, and also a major developmental signaling pathway in both lineages^31,32^ (Supplementary Figure 8A).

Interestingly, most shared terms labelled as molecular functional convergences are independent expansions observed in multicellular plants and animals, the two eukaryotic lineages hosting complex multicellular organisms (22 expansions) (Examples highlighted in Figure 3A by the red arrows and in Figure 3B, “GO” column; Figure 3C contains the treemap of the GO terms). Many of these terms are biological concepts long described as some of the key phenotypic innovations in the emergence of complex multicellularity, including components of the extracellular space/extracellular matrix, growth factors, and developmental pathways^1,33,34^ (Figure 3C; Figure 3B, GO:0007275 – *multicellular organism development*; Supplementary Figures 8 B-G).

We selected the GO term GO:0007275 to further evaluate the occurrence of functional convergences between multicellular plants and animals. In *H. sapiens*, this GO term annotates 328 genes, and a total of 261 distinct IPRs provide annotation to this GO (Supplementary Table 3). These comprise major constituents of many canonical vertebrate-specific developmental pathways, such as homeodomains, hedgehog, cadherins, Wnt, Notch, and forkhead. In *A. thaliana*, this GO annotates 161 genes, and is provided by 54 IPRs representing components of several plant-specific developmental pathways, such as EARLY FLOWERING^35^, ATNDX^36^ and CLAVATA^37^. Importantly, all but two IPR domains annotated to this GO are exclusive to either *H. sapiens* or *A. thaliana*, providing compelling evidence of molecular functional convergence (Supplementary Table 3).

Some GOs independently expanded in complex multicellular plants and vertebrates represent, to our knowledge, previously unknown molecular convergences. We highlight two GO terms where the expansion in angiosperms and vertebrates is strikingly similar, with the two lineages having much higher frequencies than any other group, including invertebrates (Figure 3B, GO:0009605 – *response to external stimulus*, GO:0048511 – *rhythmic process*).

The GO term GO:0009605 annotates 207 and 155 genes in the human and tale cress genomes, respectively (Supplementary Figure 8I, Supplementary Table 3). In the human annotation data, this GO is associated with 148 IPR IDs, while in the tale cress annotation data, 38 IPR IDs provide this association. Importantly, no IPR ID is shared across the annotations of these two genomes, therefore demonstrating the absence of homologous genes contributing for the independent expansions. The human genes annotated to this GO comprises a diverse range of molecular components to interact with the environment, such as immune system receptors (IPR000057 – *CXC chemokine receptor 2*), components of vision (IPR000378 – *Opsin red/green sensitive*) and audition (IPR030743 – *Cochlin*), appetite regulators (IPR009106 – *CART satiety factor*), and learning behavior (IPR008097 – *Fractalkine*).

The tale cress genes annotated to this GO encompasses components of the immune system (IPR022618 – *Defensin-like protein 20-27*), together with players in phenomena such as root phototropism (IPR02995 – *Root phototropism protein 2*), root and shoot response to gravitropism (IPR044683 – *LAZY family*), apoptosis (IPR031171 – *Lazarus 1*), and phytochromes with roles in seed germination, stem elongation, and organogenesis (IPR001294 – *Phytochrome*). The GO:0048511 (*rhythmic process*) annotates 11 genes in *H. sapiens* through 11 IPR IDs, while in *A. thaliana*, the same GO annotates 14 genes through 5 IPR IDs (Figure 3B, Supplementary Table 3). In both the human and the tale cress genomes, this GO annotates components of the circadian rhythm. Again, there are no shared IPR IDs among these two species. Together, these two GO terms indicate the importance of complex multicellular organisms to modify their behaviors during their life cycles, such as growth and reproduction, during their interaction with a changing environment.

### Molecular functional convergences in the evolution of metazoans

The majority of GO terms associated with NCT are observed only in a single multicellular lineage: metazoans (310 out of 352 significant associations). We observed that 11 of these GO terms also occur in the genomes of *C. owczarzaki* or *Dictyostelium discoideum*, the two closest unicellular relatives of the metazoan lineage in our analysis^38,39^, indicating a pre-metazoan origin of these biological processes. Seven terms are exclusively observed in *C. owczarzaki*, including recognizable signatures of the metazoan developmental process previously reported to be shared with this species, such components of kinase signaling pathways and cell migration processes (GO:0007169 - *transmembrane receptor protein tyrosine kinase signaling pathway*, GO:0030334 – *regulation of cell migration*, Supplementary Figures 9 A-B)^40^.

The 299 associated GO terms exclusively found in metazoans contain many biological systems and processes broadly recognized as signatures of this lineage, ranging from the cellular to the systemic level (Supplementary File 1, section “Evolution of multicellularity in metazoans”). Among others, we found expansions of crucial agents of embryogenesis, such as morphogens, growth factors, and other components of developmental signaling pathways. We also observed expansions in genes playing roles in the emergence of key biological systems of metazoans, such as skeletomuscular, digestive, circulatory and nervous, as recently reported by^41^ (Supplementary Figures 9 C-G).

## DISCUSSION

Plants and animals shared a common ancestor approximately 1.8 billions of years ago^14^. Since them, the number of cell types increased independently in these two lineages to a level of complexity not seen in other eukaryotic groups, leading to the emergence of specialized tissues, organs, and systems. A key biological question is whether this phenotypic convergence emerged due to the action of homologous elements shared from the common genomic background of eukaryotic cells or due to molecular functional convergences caused by expansions of non-homologous genes fulfilling the same biological roles.

The homologous regions associated with NCT comprise virtually metazoan- and vertebrate-specific expansions of key components for the evolution of multicellularity. However, our strategy could not find evidence of homologous regions associated with NCT that are shared across multicellular lineages and independently expanded on them. The functional-based annotation provided by GO terms, in contrast, found compelling evidence of many biological roles independently expanded in angiosperms and metazoans. As we demonstrated, these comprise molecular functional convergences of non-homologous genes long described as major players in the evolution of complex multicellularity, such as components of developmental pathways.

We also found previously unknown biological roles associated with NCT that also correspond to independent expansions of genes playing roles in the response to stimuli, immune system processes, regulatory components, and the perception of natural rhythmic processes. Our findings deepen the understanding of how complex multicellular lineages independently evolved mechanisms to coordinate the development of their distinct cell types to produce complex bodies with organs and systems, together with their genomic adaptations to interact with a changing environment.

## METHODS

### Acquiring high-quality phylogenetic, annotation and phenotypic data Estimation of number of the number of cell types (NCT)

We obtained the average Number of Cell Types (NCT) values for 56 species from the latest review available on the topic^10^. For seven species of early-branched eukaryotes we obtained cell type estimations as don by the same authors of this review: *Phytophthora infestans*, *P. sojae*, *P. parasitica* and *Achlya hypogyna*. In this case we considered their NCT values as the average value for *P. ramorum*^10^, which we used as a proxy for species within Class Oomycota, as NCT values in this taxon are mainly due to its conserved reproductive cycle^42^; *Saprolegnia parasitica* (estimated from ^43^); *Toxoplasma gondii* (estimated from^44^); *Capsaspora owczarzaki* (estimated from^45^). Information about average number of cell types and additional genomic information is available in Supplementary Table 1.

### Database of high-quality, annotated, non-redundant proteomes

We initially downloaded all genomes for the 63 species from RefSeq^46^ or GenBank^47^ databases and proceeded by extracting all protein sequence isoforms within a genome, together with locus id information, using an *in-house* software. Briefly, for each genome, we start by selecting all real coding regions (tag “translation” available in genbank files, removal of pseudogenes using tag “pseudo”). At this point, we also capture information about the locus of origin of a given protein sequence using either the “locus_tag” or “gene” tags. Such files comprise the redundant proteomes, where each locus may be represented by more than a protein sequence due events such as alternative splicing.

We proceeded by obtaining non-redundant proteomes for each species, defined as a proteome where each coding locus is represented once^11^. For that purpose we selected, for each coding locus (represented by the information stored in “locus_tag” or “gene” tags), the longest protein (for proteins of the same length, we randomly selected a sequence).

We used BUSCO^48^ to evaluate the completeness of non-redundant proteomes when evaluated against the database of 303 single-copy, almost universal orthologs shared across *Eukarya*. We used the following cutoffs to select high-quality genomes: completeness >= 70%, single copy >= 60%, duplicated genes <= 25%, missing genes <= 25%, fragmented genes <= 25%. All 46 proteomes considered suitable for downstream analyses were annotated using InterProScan^49^.

### Phylogenetic data

We found the 17 common species across these two trees to be concordant in most of their branching patterns, especially for the most recent groups (Supplementary Figure 2C). However, major differences between the trees were found in the topology of some early-diverging groups. The first discrepancy is the position of the TSAR group (represented by *Phytophthora sojae*), which is the closest group sharing a common ancestor with Archaeplastida in our list of species according to^14^.

In disagreement with current knowledge, the TToL tree considers Archaeplastida and Metazoa as a monophyletic group when considering our species list, and places the TSAR clade as a sister group of the Archaeplastida+Metazoa group. Another major difference is the position of *Dictyostelium discoideum*, one of the closest extant lineages of Opisthokonta ^50^ (represented in our analysis as the group encompassing Metazoans, *Capsaspora owczarzaki* and Fungi). The TToL phylogeny, in contrast, considers *D. discoideum* as the earliest branching species of the group encompassing all other species in our dataset.

We conclude that the tree provided by ^14^ is a suitable as a scaffold tree representing the phylogenetic branching pattern of major eukaryotic lineages, together with their divergence times. The tree provided by TToL, on the other hand, provides the correct topology and divergence times for most of the species in our dataset; more importantly, it provides phylogenetic information for the recent lineages that are not covered by the scaffold tree. We proceed by using our scaffold tree and adding the remaining 32 species to their respective positions using information from the TToL tree to get a final species tree for the 49 species (Supplementary Figure 1, “Final tree”; Supplementary Figure 2D).

### Detection of molecular functional convergences

We defined six groups of lineages based on their phylogenetic relationships and phenotypes: Protozoans, Fungi, Plant(Chlorophytes), Plant(Angiosperms), Metazoa(Invertebrates) and Metazoa(Vertebrates). As all significantly associated annotation terms that are shared across multicellular lineages are also observed in the two metazoan groups with some of the highest values, we computed the median frequencies of associated GOs for each of the six groups and defined as molecular functional convergences the GO terms where either Plant(Angiosperms) or Fungi are among the three largest non-zero median values for a given annotation term.

### Statistical analyses

We used Wilcoxon non-parametric test for all statistical tests to compare the distribution of prevalence across annotation schemas (Figure 2C). As for comparing the our results with those generated by VC^11^ under the null hypothesis of random selection of results, we computed the statistical significance of the results jointly flagged as associations with NCT using the cumulative distribution of a hypergeometric variable (for further information, see Supplementary File 1, section “Failing to take into account phylogenetic relatedness produces spurious associations”). We used CALANGO^13^, a first-principle, phylogenetically aware comparative genomics tool, to search for genotype-phenotype associations between the frequencies of annotation term and NCT values (q-value cutoff < 0.01 for the phylogeny-aware models). All figures and statistical analyses were produced using R language. All data and code used to generate this study can be obtained from Francisco Pereira Lobo.

## Abbreviations

NCT: Number of Cell Types
FDR: False Discovery Rate
GO: Gene Ontology
IPR: InterPro identifier.

## Notes

### Competing Interest Statement

The authors have declared no competing interest.

