## Supplementary File 1 for "Phylogeny-aware modeling uncovers molecular functional convergences associated with complex multicellularity in Eukarya"

#### Supplementary Results

##### Acquiring high-quality phylogenetic, annotation and phenotypic data

A crucial step when searching for functional classes of genomic components associated with a phenotypic variable is using a high-quality set of non-redundant genomic components, as biases causing variations in genomic content may mask biologically meaningful associations. A major error source are the known pitfalls in genome assembly or gene prediction procedures, which generates genome assemblies of varying qualities and makes systematic error control an imperative[1]. Other bias sources are true biological events that causes a drastic alteration in genomic content, such as the widespread secondary gene losses commonly observed in parasitic species and the whole-genome duplication events in several plant and metazoan lineages, including recent domestication events of crops[2,3].

We initially found 62 species with phenotypic (NCT values) and genomic annotation data available (see Supplementary Table 1 for a list of species and their associated metadata). After excluding the genomes with no predicted coding regions, we produced non-redundant proteomes for each species by representing each protein-coding locus by the protein sequence coded by its longest known isoform. We found the number of protein-coding genes to be within the expected values for the major groups, with protozoans, plants and metazoans having the largest median of non-redundant proteomes, and fungi having smaller proteomes[4,5] (Supplementary Figure 1B, “non-redundant proteome size”).

We proceeded by evaluating the completeness metrics as provided by BUSCO[6] (Eukaryotic dataset v10) (Supplementary Figure 1C). Among the genomes with the lowest completeness values are the fungi *Encephalitozoon cuniculi* and the protozoans *T. annulata T. gondii*, *L. major*, *T. brucei*, *P. falciparum*, and *E. histolytica*. Such organisms share some common properties: 1) a parasitic lifestyle, with considerable gene losses in important cellular processes usually found in other eukaryotes and several lineage-specific expansions of gene families associated with a parasitic phenotype [7-13]; 2) some of the smallest gene counts found in our analysis (Supplementary Figure 1C, "non-redundant proteome size"); 3) some of the highest counts of missing and fragmented orthologs (Supplementary Figure 1C, "Fragmented" and "Missing").

After considering all information, we decided to exclude these genomes from downstream analysis, since the large gene losses observed in such species are likely to be a consequence of a parasitic lifestyle, and may not truly represent the frequency of annotation terms associated with a small number of cell types. It is worth mentioning that Vogel and Chothia included the genomes of *T. brucei*, *L. major*, *P. falciparum* and *E. histolytica* in their analysis. As far as we could survey, these authors did not perform any genome quality control procedure, which was a non-canonical best practice at the time.

Some plants to have more than 25% of duplicated BUSCOs (Supplementary Figure 1C, "duplicated", 4 out of 59, ~ 7%). Plant genomes are known to be the subject of previous events of whole-genome duplication[2], especially during domestication[3]. Hence, the genomic content of these species is likely to be relatively biased due to relatively recent genomic-scale alterations and, for this reason, we excluded the species *Selaginella moellendorffii*, *Zea mays*, *Populus balsamifera* and *Physcomitrella patens* from downstream analysis.

To obtain the species tree needed to compute phylogeny-aware models, we gathered phylogenetic information from two tree sources (Supplementary Figures 1 C and E represent the scaffold and donor trees, respectively) based on our current knowledge of Eukarya evolution. We started with a scaffold tree that includes all major eukaryotic lineages included in our species list and represents the most accepted topology for major eukaryotic lineages[14] (17 species, Supplementary Figure 1C, non-marked branches correspond to shared species). A second tree with 49 out of the 50 species, including 17 shared species observed in the scaffold tree, together with their branching patterns and divergence times, was gathered from the TimeTree of life (TToL) web tool to be used as a donor tree[15] (Supplementary Figure 1E).

We found the 17 common species across these two trees to be concordant in most of their branching patterns, especially for the most recent groups (Supplementary Figure 1D). However, major differences between the trees were found in the topology of some early-diverging groups. The first discrepancy is the position of the TSAR group (represented by *Phytophthora sojae*), which is the closest group sharing a common ancestor with Archaeplastida in our list of species according to[14].

In disagreement with current knowledge, the TToL tree considers Archaeplastida and Metazoa as a monophyletic group when considering our species list, and places the TSAR clade as a sister group of the Archaeplastida+Metazoa group. Another major difference is the position of *Dictyostelium discoideum*, one of the closest extant lineages of Opisthokonta[16] (represented in our analysis as the group encompassing Metazoans, *Capsaspora owczarzaki* and Fungi). The TToL phylogeny, in contrast, considers *D. discoideum* as the earliest branching species of the group encompassing all other species in our dataset.

We conclude that the tree provided by[14] is a suitable as a scaffold tree representing the phylogenetic branching pattern of major eukaryotic lineages, together with their divergence times. The tree provided by TToL, on the other hand, provides the correct phylogenetic information for the recent lineages that are not covered by the scaffold tree. We proceed by using our scaffold tree and adding the remaining 32 species to their respective positions using information from the TToL tree to get a final species tree for the 49 species.

Importantly, this phylogenetic tree is not intended to provide the most precise reconstruction of the phylogenetic history of these groups. Instead, it is aimed at providing a reasonable phylogenetic scaffold to be used as an additional parameter in our statistical models to account for the non-independency observed in species data. The topology and the evolutionary time of major eukaryotic lineages is itself a rich and rapidly developing field, even though recent studies have started to confidently establish the evolutionary relationships of major eukaryotic groups, which was used as the scaffold tree in our analysis[16]. In fact, other phylogenetic trees can (and should) be used to investigate if the findings we report are compatible with the ones found while considering other plausible phylogenies.

Our final dataset comprises 49 eukaryotic species with high-quality non-redundant proteomes, phylogeny and NCT values available. It includes three lineages where multicellularity evolved independently: 6 fungi, 26 metazoans and 9 plants (Archaeplastida), including two lineages with complex multicellular organisms, and together with 8 early-branching unicellular/oligocellular eukaryotic species values historically referred as the parafiletic group of protists (and also referred this way from now on). These protozoan species are in key phylogenetic positions to provide a better understand of the evolution of multicellularity in Opisthokonta (the clade including Fungi and Metazoa), Metazoans and plants.

**NCT value of *Homo sapiens* is biased when compared with other species and influences downstream analyses**

The most recent average NCT value reported for *Homo sapiens* used in a comparative genomics study is 264.5[17]. This value is highly discrepant from the other mammals, which have arguably the same overall degree of complexity (e.g. the NCT value for the chimpanzee *Pan paniscus* is 169, resulting in a NCT value 56.5% greater for *H. sapiens*). The discrepancy is likely to be due to an updated review of the number of cell types for our species, which described a total 411 cell types and inflated the average counts obtained in[18]. This number is likely to further increase in the near future with the increasing number of studies of large-scale molecular characterization of the human body at the single-cell level[19].

The large amount of single-cell data recently produced for *H. sapiens* allows the subtle characterization of many cell (sub)populations, which may arguably be named distinct cell types, and certainly inflated NCT values for our species. The ongoing effort to provide standardized single-cell characterization of model organisms by using controlled dictionaries of cell types in a phylogeny-aware statistical scaffold is expected to fill this gap in the coming years, as has recently done for metazoans[20]. However, the current lack of deepness in the phenotypic characterization of cell types in non-human species is likely to be the reason for the discrepancy in the average NCT value for *H. sapiens* and other species.

By using the homologous regions and GO terms as defined in the distinct annotation schemas, we used three experimental designs to evaluate how the human NCT value could bias downstream analyses: 1) a standard analysis using 264.5 as NCT value for *H. sapiens*; 2) the removal of *H. sapiens* from the species dataset; 3) using the NCT value for *P. paniscus* (169) as a proxy for *H. sapiens*.

The experiment using the original NCT value for *H. sapiens* produces results that are highly discrepant when compared with either the removal of *H. sapiens* from our set of species or using the NCT value for *P. paniscus* as a proxy for *H. sapiens* (Supplementary Figure 2). Furthermore, the removal of *H. sapiens* and the NCT proxy strategy produce highly concordant results, as they share the vast majority of associated terms. We decided to proceed using the "*Pan* as proxy" phenotype parameters, as this allowed us to include arguably one of the best mammalian genome assembly available while also considering the NCT bias in *H. sapiens*. Also for this reason, the NCT value depicted in Figure 1 for *H. sapiens* is 169.

**Analysis of homologous regions associated with NCT and broader taxonomic distribution**

We found three associated homologous regions that occur exclusively in protozoans and metazoans. The first region is IPR039008 (*Intermediate filament, rod domain*) which was observed in all metazoans (with the exception of the Placozoa *Trichoplax adhaerens*), in *C. owczarzaki*, the closest unicellular relative of Metazoans in our dataset, and in three out of the four protozoans from the TSAR group (the sister group of Archaeaplastida in our dataset) (Figure 3B). Intermediate filaments are traditionally considered a large gene family coding key components of the metazoan cytoskeleton, with major roles in determining cell shape diversity long thought to be metazoan-specific[21]. More recently, homologous of intermediate filaments components of the nuclear membrane were described in *D. discoideum*, *C. owczarzaki* and in the TSAR group, together with an absence in the basal metazoan *T. adhaerens* and in plants and fungi, which suggests an ancient eukaryotic origin followed by secondary losses in plants, fungi and at least one metazoan lineage[22]. This is the same pattern found in our analysis, providing evidence that our automatic annotation schema is capturing the phylogenetic distribution profile of homologous regions associated with cell diversity. Together, we conclude that the expansion of this homologous region in species with higher NCT values appear to be metazoan-restricted.

The second case is *IPR018487 - Hemopexin-like repeats*, observed in all metazoans (with the exceptions of the sponge *Amphimedon queenslandica* and of *T. adhaerens*) and in *C. owczarzaki* (Supplementary Figure 3B). The hemopexin-like fold is an ancient, variable and widespread protein fold found in bacteria, virus and eukaryotic species, and represented by several IPR IDs[23,24]. As we could not find any previous works describing the systematic phylogenetic distribution of this domain, we proceeded by evaluating the taxonomic prevalence of the hemopexin IPR ID associated with NCT (IPR018487) in the Interpro database. As of December 2023, This domain occurs in 1,854 proteomes, including viruses (10 species), prokaryotes (459 bacteria and 5 archaea species) and eukaryotes (1380 species).

The vast majority of sequences containing this domain come from metazoans species, with an occurrence in early branching eukaryotes that is compatible with our findings (presence in *C. owczarzaki* and absent from TSAR and from *D. discoideum*). Importantly, even though this domain is observed in several plant and fungi species, it is far from being universally observed in these lineages. When considering the plant species in our analysis, this domain is observed only in two protein sequences of the proteome of *Oryza sativa* as provided by the InterPro database.

As we have not detected any hemopexin in our non-redundant proteome for *O. sativa*, we used the sequence described from one of these sequences (GenBank ID: CAE03710) as a protein query to search the entire predicted redundant proteome described in the genome assembly used in our analysis using blast, but fail to find any similar sequences. A search using tblastx (protein query against the translated genomic sequence) with the same protein sequence, however, found an identical region in the genome assembly used in our analysis that, however, is not annotated as a protein-coding gene. Additional searches using blastp in other genome assemblies from the genus *Oryza* found highly similar proteins annotated as protein-coding genes in four assemblies out of the 98 currently available as of December 2023 in NCBI Genome database. We conclude that this sequence is not predicted as protein-coding gene in the genome of *O. sativa* used in our analysis and in most other assemblies publicly available. The extension of our search using blastp found similar sequences to be spread across angiosperms, but truly absent from at least in the predicted coding sequences from species belonging to all the genera of plants in our dataset.

Proteins with hemopexin-like fold are poorly characterized in plants, and appear to be involved in different cellular pathways. In the grass pea *Lathyrus sativus*, a protein with hemopexin-type fold was described as a sensor of oxidative stress through a reversive heme binding[25]. Other examples of a hemopexin-like fold in plants are found in seed storage proteins and lectins[26,27]. More recently, one of the two coding sequences of hemopexin in *O. sativa* has been characterized as a structurally novel hemopexin fold playing a role in chlorophyll degradation[28].

In fungi, our manual search for protein entries containing IPR018487 in the InterPro database found it to be present in hypothetical protein sequences in genomes from the genus *Aspergillus*, even though absent from the genomic sequence of the species used in our analysis (*A. nidulans*). We used one of the sequences from this genus containing the IPR ID as query (genbank ID: PYI13027.1 from *A. violaceofuscus*) for a blastp search against the full nr database. We found this protein to be restricted to hypothetical sequences from Fungi, mostly observed in this genus, and also in the Hepatophyta *Marchantia paleacea*. This domain is not observed in any other genera of fungi included in our dataset. A thorough literature review found this domain to be poorly characterized in fungi[29].

This domain is a telltale of how noisy genomic data can be, even when starting with high-quality genome assemblies. To affirm that IPR018487 is completely absent from plants and fungi is certainly not true, as demonstrated in our searches in sequence databases and in our literature review. However, our strategy of using a phylogenetically diverse set of high-quality proteome sets, produced by distinct research group is also a strong manner to prevent that any systematic bias in these procedures affects our general conclusions. Even though lacking a more profound characterization in both plants and fungi, genes coding for this domain appear to have roles that may contribute to the emergence of multicellularity at least in plants. However, this homologous region lacks the phylogenetic broadness to be confidently claimed as a major factor contributing for the independent emergence of multicellularity in Eukarya. Our extensive sequence search using blast found it to be truly absent from all the genomes of plant and fungi used in our analysis (with the special case of *O. sativa* described above), even though these are complex eukaryotic lineages.

In vertebrates, hemopexin domains are found in a family of evolutionary related proteins, including a serum glycoprotein that binds and transports haem to the liver for breakdown and a heavily expanded group of genes coding for matrix metalloproteinases that play critical roles in a wide range of cell behaviors such as proliferation, migration and differentiation[23]. Further inspection of the two *C. owczarzaki* proteins containing the hemopexin domain found them to also contain putative DNA binding domains, a role that, as far as we are aware, is not shared by any of the metazoan proteins where the hemopexin domain is also observed.

The presence of two copies of genes coding for hemopexin domains, together with the absence of sequences coding for this domain in *A. queenslandica* and of *T. adhaerens*, suggests a secondary loss in these metazoan lineages. However, the broad phylogenetic distribution of this domain, ranging from viruses to all cellular lineages, also suggests that its origin may predate the emergence of cellular life. The absence of this domain in several lineages of bacterial, fungi and plants also suggests that this domain may have been recurrently lost, horizontally transferred across lineages, or that its sequence has diverged beyond recognition by our protocol. Together, we conclude that the phylogenetic distribution of this domain does not present solid evidence of the independent expansions in multicellular lineages associated with NCT, even though it has a clear signature of a vertebrate-specific expansion. Further characterization of the two genes of *C. owczarzaki* coding for this domain may provide additional information about the contribution of this homologous region for the evolution of multicellularity in metazoans.

The third and last homologous region associated with NCT also observed in protozoans and metazoan species is a ubiquitin-associated protein (UBAC), observed in *C. owczarzaki* and all metazoans but the basal lineages *A. queenslandica*, *T. adherens*, *Hydra vulgaris* and the two roundworms (*Caenorhabditis elegans* and *C. briggsae*) IPR022166 (UBAP2/protein lingerer). The taxonomic distribution of this domain in the InterPro database largely support our observations: in *D. melanogaster*, this domain is found once in the single-copy gene *lingerer*, which is a component of a major negative regulator of tissue growth[30]. The two copies of this domain in *H. sapiens* code for the genes UBAP2 and UBAP2L, and have been recently characterized as the orthologs of yeast gene *Def1*, a component of the transcription-coupled nucleotide excision repair pathway[31]. However, the protein sequence of the yeast Def1 protein lacks this homologous region, suggesting that additional metazoan-specific roles for these genes cand be provided by this domain. Interestingly, both UBAP2 and UBAP2L are overexpressed in multiple types of cancer[32]. This observation, together with the biological role of the *D. melanogaster lingerer* gene, suggests that this domain may play key roles in tissue growth in humans not fully described.

**Evolution of multicellularity in metazoans**

**Signature system of vertebrates**

Metazoans have a set long recognized systems that are signature traits of this lineage, including nervous, muscular, circulatory, digestive and endocrine. We found expansions of genes annotated to GO terms that represent all these systems (Supplementary Figures 9C-F). We also observed an expansion of genes coding for collagens (GO:0062023 - *collagen-containing extracellular matrix*, Supplementary Figure 9L). Both GOs represent metazoan-specific components of connective tissues already described as expanded in chordates when compared to non-chordate metazoans[33,34].

**Morphogens and growth signaling pathways**

We found GO terms associated with NCT occurring exclusively in vertebrates that describe distinct components of classic developmental signaling pathways known to have critical roles in metazoan embryogenesis: morphogens and growth factors. Morphogens are signaling molecules that coordinate cell fate during embryonic development through gradient variation within cells, tissues and body plans, establishing some of the long-range communication needed for proper development of body plans, tissues, organs and systems, and growth factor are ligands that activate signaling pathways that stimulate cell growth, proliferation and differentiation[36,38]. Among the morphogens we highlight the vascular endothelial growth factor (GO:0048010), a known morphogen during kidney embryogenesis[37], and the retinoic acid (GO:0005501), a morphogen that controls the differentiation and patterning of various stem/progenitor cell populations during the chordate developmental program[40] (Supplemenary Figures 9 I-J).

Growth factor and their receptors is an umbrella term describing a heterogeneous set of protein and peptide ligands and their receptor pairs that activate many signaling pathways. These were named after such ligands, and with major roles in the development and homeostasis of metazoan-specific tissues, organs and systems, such as skin, connective tissue, blood, muscle and bone[38,46,47]. We found expansion patterns in several of these growth factor signaling pathways and their components, such as fibroblast growth factor (FGF), vascular endothelial growth factor (VEGF, GO:0005021), platelet-derived growth factor (PDGF, GO:0048008), and neurotrophins (GO:0005030). As ana example, FGFs are a group of homologous genes coding for signaling polypeptide hormones essential during embryonic development of epithelium, bone and soft connective tissues, being also homeostatic factors in injury response and metabolic changes. In *H. sapiens,* the FGF gene family comprises 22 paralogs, and these have already been described as expanded in chordate lineages[38]. We found both GO terms describing the pathway as a whole as expanded in chordates (GO:0008543 - *fibroblast growth factor receptor signaling pathway*), as well as specific components of this pathway, specifically the FGF ligands (GO:0005104 - *fibroblast growth factor receptor binding*) (Supplementary Figures 9F-G).

**Failing to take into account phylogenetic relatedness produces spurious associations**

The work by Vogel and Chothia (VC dataset/work) is a landmark study in integrating phenotypic and genomic data to study the evolution of multicellularity in Eukarya. These authors used the SUPERFAMILY database, together with NCT values for 38 eukaryotic lineages, to search for annotation IDs associated with NCT values (defined by them as the ones with Pearson's correlation values either > 0.8 or < -0.8). They found a strong bias towards vertebrate-specific homologous regions associated with NCT, together with an enrichment of extracellular processes, signal transduction and components of the immune system[49].

However, as previously mentioned, the methodology used by these authors do not consider two major factors, undoubtedly because these were not established protocols in large-scale comparative genomic analyses as they are nowadays: 1) account for the non-independence of species data due to common ancestry, 2) the inclusion of several species with low BUSCO values due to specific evolutionary events that are likely to bias relative term frequency (e.g. recent whole-genome duplications in plants and massive gene losses in parasites). Therefore, even though the VC work is a pioneering study for the understanding of the genomic evolution of complexity in Eukarya, it also contains major issues that may bias its conclusions and deserve further investigation.

Our dataset vary from the one used by VC in major aspects, such as species numbers (38 in VC and 49 in our dataset, with an intersection of 25 species), the estimation of NCT values, which were updated for several species since their work, and the representation of the abundance of annotation terms (relative frequencies of annotation terms within each genome in our case, and the relative abundance of a SUPERFAMILY term in relation to its abundance in other genomes, in the case of the VC dataset). Therefore, we started our analysis by evaluating whether our initial datasets and data pre-treatment procedures produce sufficiently similar results when compared with the work of VC when using the same statistics. This analysis allows us to known if any solid conclusions can be drawn while comparing our results with the ones found by VC.

The VC dataset comprises 1,219 SUPERFAMILY annotation terms, of which 194 were deemed as associated with NCT values. From the set of 1,458 SUPERFAMILY annotation terms in our annotation data, a total of 1,164 are shared with the VC dataset. For those, we used the same statistics and cutoffs as VC (Pearson's correlation >= 0.8 or <= -0.8) to emulate their study as much as possible. A total of 82 homologous regions were found to be associated with NCT in our dataset, 62 of which were also flagged by VC in the shared SUPERFAMILY annotation terms between our analyses. To test the concordance of the significant correlations found by us and those reported by VC, we performed a test of hypothesis assuming the VC results as fixed and verifying the significance of the number of concordant results we found. In brief, VC reported K = 194 significant correlations out of N = 1,164 shared possibilities, and we found n = 82 significant results, out of which 62 were in agreement with those suggested by VC.

Under the null hypothesis of random selection of correlations as significant by us, the statistical significance of these results can be calculated based on the cumulative distribution of a hypergeometric variable (CDF: Cumulative Distribution Function; Population size: N = 1,164; Number of success states: K = 194; Sample size: n = 82):

**p = 1 - CDF(N=1164, K = 194, n = 82, k = 64)**

Which yields an approximate p-value of 1.68e-38. This finding strongly suggests that our dataset shows a remarkable concordance with the VC data in terms of which correlations it flags as relevant. Therefore, we consider our annotation and phenotypic data to be sufficiently similar to the original VC data, which allows further investigation of how the phylogeny-aware statistical models and multiple hypothesis correction used in our analysis impact the results.

However, we found none of the distinct 1,458 homologous region represented in SUPERFAMILY to be significant associated with NCT values when using phylogeny-aware models (q-value < 0.01). We concluded that using traditional statistics (e.g. Pearson's correlation) to search for associations between the relative abundance of homologous regions and NCT values produces spurious associations that fail to be observed in the same dataset when phylogenetic information is taken into account.

### SUPPLEMENTARY FIGURE LEGENDS

**SUPPLEMENTARY FIGURE 1 – Generation of annotation and phylogenetic data.** A) Definition of the annotation schemas. In this picture we represent three hypothetical proteins (PROT_001-PROT_003) annotated to either a function-based (GO-based annotation) or a homology-based annotation schema (Homology-based annotation). The three proteins are annotated to four distinct homologous regions (IPR boxes in orange, brown, purple and green). These four homologous regions are associated with two GO IDs (black boxes). Each GO ID annotates non-homologous regions that fulfill distinct roles (GO:0035480 – orange and green boxes, GO:0007049 – purple and brown boxes). Genomes are annotated by generating unique pairs of proteins (Protein_ID) and annotation terms (Annotation_ID) for each genome, followed by the computation of the sum of all annotation terms from the pairs in a given genome ("Sum" column) and the computation of relative frequencies of annotation terms ("Frequency" colum), defined as the ratio of the sum of occurrences of an annotation term in a genome and the sum of occurrences of all annotation terms in the same genome. The annotation of non-homologous regions that fulfill the same biological role to the same GO ID increases the prevalence of these terms in comparison with the noes provided by homology-based schemas ("Frequency" column). B) Genome quality metrics. "Non-redundant proteome size" – number of protein-coding genes for each genome, represented as the longest known isoform of each locus; The remaining plots represent BUSCO results for Completeness, Single-copy, Duplicated, fragmented and missing groups. Species acronyms are the same as in Figure 1B. C) Scaffold tree to build the final phylogenetic tree. Shaded taxa denote excluded branches. The remaining groups were used as anchors to stich the branches from the donor tree. D) Comparison of scaffold and donor tree. Inconsistencies were found for the TSAR group (*Phytophthora sojae*, P_soj) and for *Dictyostelium discoideum* (D_dis). Species acronyms are the same as in Figure 1B. E) Donor tree from TimeTreeOfLife. Shaded taxa denote groups to be included in the anchor branches of the scaffold tree (non-shaded groups in this tree and in tree of Supplementary Figure 1C).

**SUPPLEMENTARY FIGURE 2 – Comparison of experiments using distinct NCT values for *H. sapiens*.** A) Venn diagrams of homologous regions associated with NCT in the three experiments to investigate how the larger NCT value for *H sapiens* could be biasing our downstream analyses. less_H_sapiens (excluding *H. sapiens* from our data); Pan_as_proxy (using *P. troglodytes* NCT values as a proxy for *H. sapiens*); orginal_NCT_value (using the average NCT data available from the literature). B) A) Venn diagrams of GO terms associated with NCT in the three experiments to investigate how the larger NCT value for *H sapiens* could be biasing our downstream analyses. less_H_sapiens (excluding *H. sapiens* from our data); Pan_as_proxy (using *P. troglodytes* NCT values as a proxy for *H. sapiens*); orginal_NCT_value (using the average NCT data available from the literature).

**SUPPLEMENTARY FIGURE 3 – Examples of homologous regions associated with NCT values and observed in metazoans and protozoans.** Left: association between phylogenetically independent contrasts (PIC) for NCT and for the relative frequency of annotation terms. The blue line represents the linear model of PICs; Correlation values are Pearson's correlation of PIC values; q-values are corrected p-values for the linear models. Right: counts and relative frequency of the annotation terms for the six groups investigated in this analysis. Each violin plot represents the distribution of values for the group; black bars are the medians of group.

**SUPPLEMENTARY FIGURE 4 – Examples of homologous regions associated with NCT values representing key components of multicellularity in metazoans (extracellular matrix, ion channels, signaling pathways, epigenetic regulators and caspases).** Left: association between phylogenetically independent contrasts (PIC) for NCT and for the relative frequency of annotation terms. The blue line represents the linear model of PICs; Correlation values are Pearson's correlation of PIC values; q-values are corrected p-values for the linear models. Right: counts and relative frequencies of the annotation terms for the six groups investigated in this analysis. Each violin plot represents the distribution of values for the group; black bars are the medians of group.

**SUPPLEMENTARY FIGURE 5 – Examples of homologous regions associated with NCT values representing key components of multicellularity in metazoans (developmental pathways) and poorly characterized gene families.** Left: association between phylogenetically independent contrasts (PIC) for NCT and for the relative frequency of annotation terms. The blue line represents the linear model of PICs; Correlation values are Pearson's correlation of

PIC values; q-values are corrected p-values for the linear models. Right: counts and relative frequencies of the annotation terms for the six groups investigated in this analysis. Each violin plot represents the distribution of values for the group; black bars are the medians of group.

**SUPPLEMENTARY FIGURE 6 – Examples of homologous regions associated with NCT values representing key components of multicellularity in vertebrates and poorly characterized gene families.** Left: association between phylogenetically independent contrasts (PIC) for NCT and for the relative frequency of annotation terms. The blue line represents the linear model of PICs; Correlation values are Pearson's correlation of PIC values; q-values are corrected p-values for the linear models. Right: counts and relative frequencies of the annotation terms for the six groups investigated in this analysis. Each violin plot represents the distribution of values for the group; black bars are the medians of group.

**SUPPLEMENTARY FIGURE 7 – Examples of GO terms associated with NCT values representing key biological functions for the emergence of multicellularity in metazoans.** Left: association between phylogenetically independent contrasts (PIC) for NCT and for the relative frequency of annotation terms. The blue line represents the linear model of PICs; Correlation values are Pearson's correlation of PIC values; q-values are corrected p-values for the linear models. Right: counts and relative frequencies of the annotation terms for the six groups investigated in this analysis. Each violin plot represents the distribution of values for the group; black bars are the medians of group.

**SUPPLEMENTARY FIGURE 8 – Examples of GO terms associated with NCT values representing key biological functions for the independent emergence of multicellularity in Eukarya.** Left: association between phylogenetically independent contrasts (PIC) for NCT and for the relative frequency of annotation terms. The blue line represents the linear model of PICs; Correlation values are Pearson's correlation of PIC values; q-values are corrected p-values for the linear models. Right: counts and relative frequencies of the annotation terms for the six groups investigated in this analysis. Each violin plot represents the distribution of values for the group; black bars are the medians of group.

**SUPPLEMENTARY FIGURE 9 – Examples of GO terms associated with NCT values representing key biological functions for the emergence of multicellularity exclusively observed in Metazoans.** Left: association between phylogenetically independent contrasts (PIC) for NCT and for the relative frequency of annotation terms. The blue line represents the linear model of PICs; Correlation values are Pearson's correlation of PIC values; q-values are corrected p-values for the linear models. Right: counts and relative frequencies of the annotation terms for the six groups investigated in this analysis. Each violin plot represents the distribution of values for the group; black bars are the medians of group.
